# The epigenomic landscape of bronchial epithelial cells reveals the establishment of trained immunity

**DOI:** 10.1101/2025.02.03.636250

**Authors:** Jeanne Bigot, Rachel Legendre, Juliette Hamroune, Sébastien Jacques, Mathieu Legars, Loïc Guillot, Harriet Corvol, Christophe Hennequin, Juliette Guitard, Jean-Yves Coppée, Viviane Balloy, Claudia Chica

**Affiliations:** Sorbonne Université, Inserm, Centre de Recherche Saint-Antoine, CRSA, AP-HP, Hôpital Saint-Antoine, Service de Parasitologie-Mycologie, F-75012 Paris, France; Institut Pasteur, Université Paris Cité, Bioinformatics and Biostatistics Hub, F-75015 Paris, France; Institut Cochin U1016, INSERM UMR8104 CNRS, 24, rue du Fg St Jacques, Paris, France; Gritstone bio, Inc., Emeryville, CA, USA; Sorbonne Université, Inserm, Centre de Recherche Saint-Antoine, Paris, France; Sorbonne Université, Inserm, Centre de Recherche Saint-Antoine, CRSA, AP-HP, Hôpital Trousseau, Service de Pneumologie Pédiatrique, F-75012 Paris, France; Institut Pasteur, Université Paris Cité, Plate-forme Technologique Biomics, Paris, France

**Keywords:** Innate immune memory, bronchial epithelial cells, epigenetic, transcriptomic, self-organizing maps, trained immunity, gene expression, joint epigenome and transcriptome analysis, quantitative regulation, transcription factor footprinting

## Abstract

**Background:** Innate immune memory, also called trained immunity, refers to the ability of innate immune cells to gain memory characteristics after transient stimulation, resulting in a nonspecific modified inflammatory response upon secondary remote challenge. Bronchial epithelial cells (BECs) participate in innate immune defence and are the first cells of the lower respiratory tract to encounter inhaled pathogens. We recently showed that BECs are capable of innate immune memory after preexposure to *Pseudomonas aeruginosa* flagellin through epigenetic mechanisms. In the present study, we investigated such mechanisms through the modification of chromatin architecture induced by flagellin preexposure that results in subsequent changes of gene expression.

**Results:** By conducting an unsupervised approach to jointly analyse chromatin accessibility and gene expression, we mapped the remodelling of the epigenomic and transcriptomic profiles during the establishment of BECs memory. We identified a Memory regulatory profile induced by flagellin exposure. It includes clusters of upregulated genes related to inflammation that are linked to a sustainable gain in chromatin accessibility and with an increased activity of specific factors (TFs) whose binding may drive this process.

**Conclusions:** In summary, we demonstrated that flagellin exposure induced changes in chromatin condensation in BECs, which sustains the reprogramming of transcriptional patterns

## Background

Host immunity can be categorised into innate and adaptive immunity. Until the 2010s, the innate immune response was described as devoid of memory capacity, unlike adaptive immunity. Innate immune cells can recall their first contact with a microorganism to remotely modulate their responses when exposed to secondary microbial encounters (1). This process, called innate immune memory or trained immunity, can be induced by the interaction of cellular pattern recognition receptors (PRRs), such as Toll-like receptors (TLRs), C-type lectin receptors (CLRs), retinoic acid-inducible gene-I-like receptors (RLRs), and NOD-like receptors (NLRs), with specific microbial molecular motifs known as pathogen-associated molecular patterns (PAMPs), leading to specific activation of immune cells. The main characteristic of innate immune memory is the maintenance of the homeostatic state induced by a first stimulus, which results in increased (training) or decreased (tolerance) cellular responses to secondary challenges. These modifications are induced by long-term epigenetic reprogramming, such as chromatin remodelling, gene transcription changes, and profound metabolic changes, leading to a nonspecific immune response to subsequent challenges (1,2).

To date, innate immune memory has been attributed to haematopoietic lineage cells, such as macrophages/monocytes (3,4), natural killer cells (5,6), and dendritic cells (7), which are major cells related to innate immunity. Nonetheless, studies have shown that innate immune memory can be carried by nonmyeloid cells, such as endothelial and epithelial cells (8). Epithelial cells, which act as barriers of the skin, airways, and intestines, form essential interfaces and detect and respond to different signals, such as innate immune cells (8). It has been demonstrated that immune memory mediated via long-term epigenetic changes can be acquired by respiratory epithelial progenitor cells during allergic inflammatory disease (9). Respiratory epithelial cells and, more specifically, bronchial epithelial cells (BECs) constitute the initial physical barrier against inhaled microorganisms. BECs express PRRs, allowing these sensors to sense and discriminate pathogens through the recognition of PAMPs, resulting in the production of antimicrobial compounds and the synthesis of inflammatory mediators. This is the case for interleukin-8, a potent neutrophil attractant and activator that participates in pathogen clearing (10). Functional defects in some chronic respiratory pathologies, such as cystic fibrosis or chronic obstructive pulmonary disease, favour bronchial colonisation by several pathogens, such as *Staphylococcus aureus*, *Pseudomonas aeruginosa* and *Aspergillus fumigatus* (11), resulting in excessive inflammation associated with a progressive decline in respiratory function.

In a recent study, we demonstrated that memory acquisition by BECs after previous contact with a bacterial compound induced nonspecific immunomodulation of their responses to subsequent reinfection (12,13). We found that BEC preexposure to *P. aeruginosa* flagellin could increase the inflammatory response to a second stimulus induced by a live fungal pathogen, *A. fumigatus,* or, conversely, decrease the inflammatory response against bacterial lipopolysaccharide stimulation. There are multiple and reversible epigenetic mechanisms that underlie innate immune memory. We previously reported that histone modifications are involved in BEC innate immune memory, suggesting the involvement of epigenetic mechanisms such as changes in chromatin states (12). Our current aim was to demonstrate that flagellin exposure induced changes in chromatin condensation in BECs, which sustains the reprogramming of transcriptional patterns. To obtain a joint quantitative description of the chromatin condensation and gene expression landscape underlying the establishment of innate memory, we combined an assay for transposase-accessible chromatin with high-throughput sequencing (ATAC-seq) technology and a microarray using DNA and RNA extracted from BECs exposed or unexposed to flagellin. We then summarised chromatin accessibility and gene expression heterogeneity by applying self-organising maps, an unsupervised machine learning technique that produces a two-dimensional representation of any high-dimensional dataset while preserving its underlying structure (14–17). By integrating accessibility and expression dynamics, we describe the different regulatory profiles underlying the BEC response to flagellin exposure. Through this joint analysis, where the transcriptome guides the classification of the epigenome, we identified a specific set of regions that undergo a persistent increase in chromatin accessibility upon flagellin exposure that is correlated with (i) the steady upregulation of associated immune inflammatory pathways over time and (ii) a greater binding probability of a distinct subset of transcription factors. Our results confirm the reprogramming and memorisation abilities of BECs through chromatin state modulation.

## Results

Using the same experimental procedures as our previous study (12), we evaluated modifications in chromatin accessibility specifically induced by flagellin pre-stimulation, which persist over time during epithelial cell proliferation and impact the transcriptome (**Figure 1A**). First, we validated a trained response after BEC exposure to flagellin in our experimental conditions by observing a significant increase in IL-8 and IL-6 synthesis after *A. fumigatus* infection at Day (D)6 compared to flagellin-naïve BECs (**Figure 1B**).

**Figure 1:**
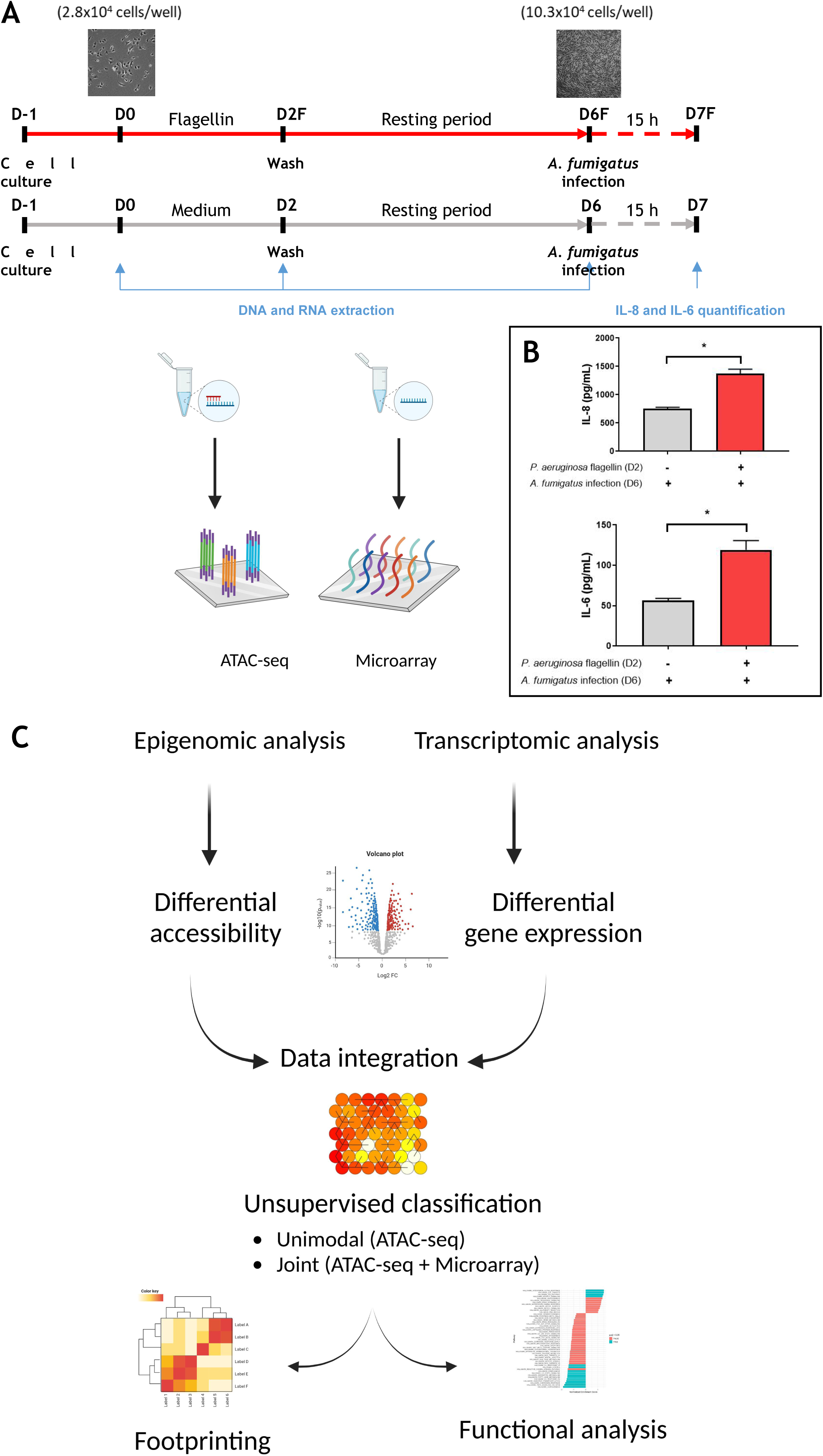
Experimental design to study epigenomic and transcriptomic modulation upon flagellin priming in bronchial epithelial cells. **A.** On day −1 (D-1), BEAS-2B cells were cultured in 96-well plates (1.5 × 10^4^ cells/well). On day (D0), the cells (2.8 × 10^4^ cells/well) were prestimulated (F) (red arrow) or not (Ctl) (gray arrow) with 5 µg/mL ultrapure flagellin (InvivoGen) from *Pseudomonas aeruginosa* for 48 h. The cells were then washed twice with media on day 2 (D2 and D2F), followed by a resting period of 4 days, until they reached confluence (10.3 × 10^4^ cells/well) on day 6 (D6 and D6F). RNA and DNA were extracted at days 0, 2 and 6 from BECs exposed (F) or not exposed (Ctl) to flagellin and then analysed via microarray and ATAC sequencing. In parallel experiments, to validate the trained response, BECs prestimulated with or without flagellin were infected with *A. fumigatus* (MOI of 1) for 15 h (dotted lines) on day 6. **B.** Supernatants were collected to quantify IL-8 and IL-6 levels by ELISA. The results are expressed after subtraction of values obtained for unstimulated conditions. Each histogram represents the mean ± SEM of triplicate samples; n=3; * p<0.05 (Mann‒Whitney test). **C.** Transcriptomic and epigenomic dynamics were analysed by differential analysis between days 2 and 6 *vs.* day 0 in BECs exposed to flagellin (F) or not exposed (Ctl). Accessible regions and genes undergoing significant expression changes were then jointly classified using an unsupervised machine learning approach to identify clusters with common regulatory profiles. The latter were functionally characterised by gene set enrichment analysis and transcription factor footprinting. D: day, F: *P. aeruginosa* flagellin, Ctl: control. Created with BioRender.com

### Flagellin exposure induces sustained chromatin accessibility changes

We used ATAC-seq to profile chromatin accessibility in the BEC genome upon flagellin stimulation (**Figure 1C**). We performed a differential analysis of accessibility changes in BECs trained or not trained by flagellin exposure during the time course, comparing D2 *vs* D0 and D6 *vs* D0. We found 29861 differentially accessible (DA) peaks (adjusted p value < 0.05), 16216 of which changed specifically in each comparison and 13645 of which changed in two or more comparisons. Interestingly, 78% (23389/29861) of the DA peaks occurred only at the end of the time course: 10108 only upon F (flagellin) exposure, 6796 only in the Ctl (control) condition, and 12957 in both. Notably, there were 608 peaks whose change in accessibility was significant at both D2F and D6F (**Additional file 1: Fig. S1A**). These results are consistent with the principal component analysis (**Additional file 1: Fig. S1B**), which shows that the primary source of variability of chromatin accessibility coincides with the difference between D0/D2 and D6 (PC1: ∼40%), followed by the difference between the Ctl and F conditions (PC2: ∼15%). Taken together, these data show that exposure to flagellin induces changes in chromatin accessibility that are absent in the unexposed condition. Additionally, for some regions, such changes were observed at D2F and were maintained up to D6F, suggesting the establishment of epigenetic memory.

### Flagellin exposure induces specific gene regulation

To complete the description of our system, we conducted differential gene expression analyses using microarray data collected from BECs exposed or unexposed to flagellin (**Figure 1C**). We observed that 4502 genes had significant expression changes in all comparisons (adjusted pval < 0.05, absolute fold change > 1). However, the expression of 1308 and 1891 genes changed only in the Ctl and F groups, respectively (**Additional file 1: Fig. S1C**). Unlike chromatin accessibility, the primary source of variability in gene expression coincided with the difference between D0 and D2/D6 (PC1: ∼70%), followed by the difference between D2 and D6 (PC2: ∼11%), indicating that major changes in gene expression were already present in the early stages of the Ctl and F time courses (**Additional file 1: Fig. S1D**). The final source of variability is the difference between the Ctl and F conditions (PC3: ∼5%). Notably, there is a global differential effect of flagellin at D2F and D6F, suggesting the establishment of flagellin-specific memory at D2F and maintenance until D6F in some transcriptomic programs specific to the F time course.

### Unimodal profiles of chromatin accessibility are insufficient to discriminate gene expression changes

To test the correlation between these sustained changes in expression upon flagellin stimulation and changes in chromatin accessibility, we used the epigenomic and transcriptomic data to identify and functionally characterise the underlying the regulatory links (**Figure 1C**).

We first characterised chromatin accessibility changes over time and identified differences between the Ctl and F conditions. We performed unsupervised classification using self-organising maps (SOMs) and clustering with the k-means approach. For this analysis, we focused on 29861 dynamic regions showing significant changes in accessibility between the Ctl and F. For regions gaining accessibility, we identified ten clusters on the resulting SOM that included three profiles: (i) one showing a sustained gain in accessibility stronger under F than under the Ctl (clusters 1, 2, 3, 5, 6, and 8); (ii) a profile with more important changes only for Ctl (clusters 4, 9, and 10); (iii) a profile with similar changes in F and Ctl (cluster 7) (**Figure 2A**).

**Figure 2:**
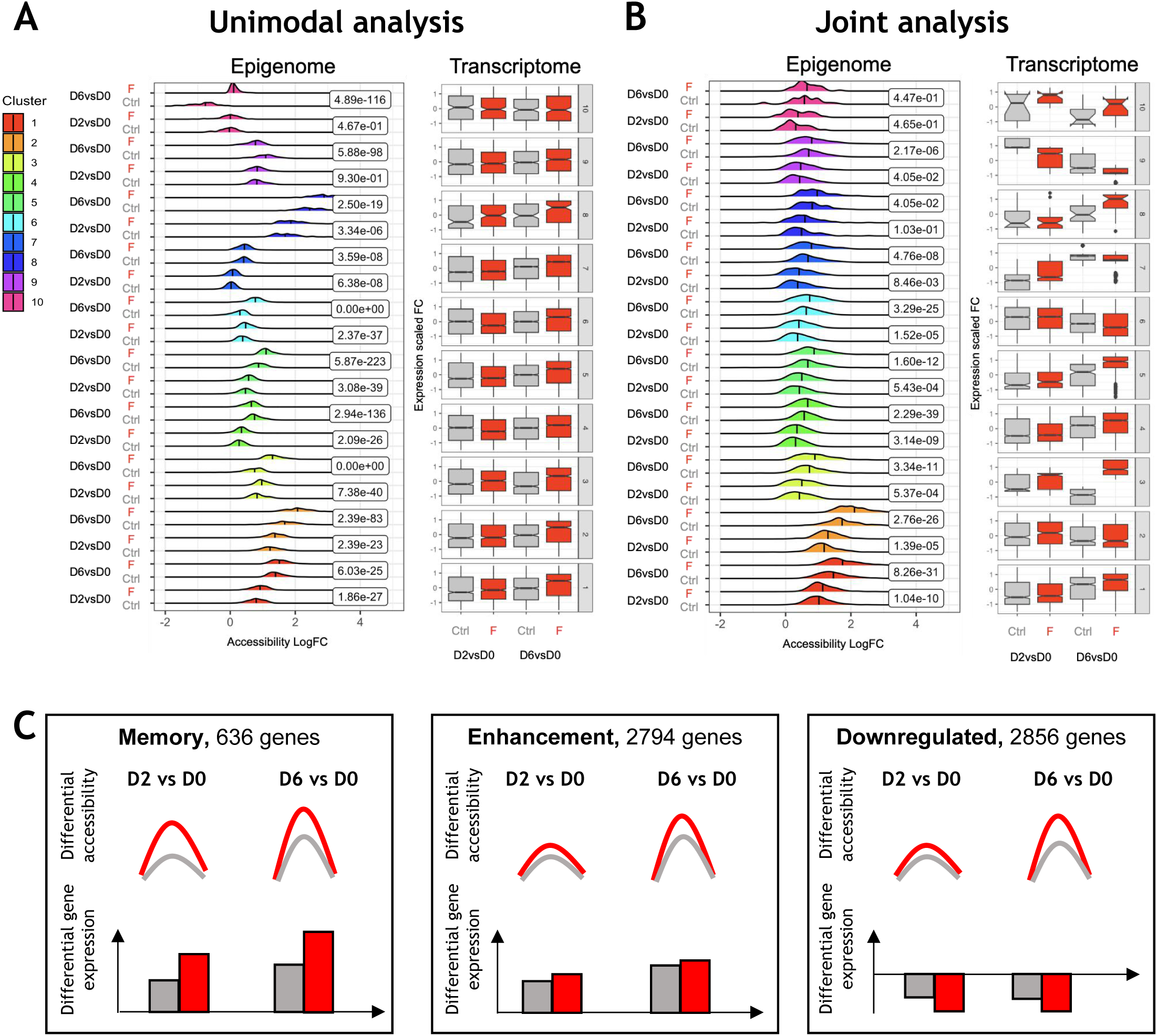
Changes in the epigenome and transcriptome during BEC priming. The data were summarised using the self-organising map (SOM) algorithm and clustered via the k-means method. Classification was performed using only accessibility data (unimodal analysis **A**) and both accessibility and expression data (joint analysis **B**). **A-B** Regions showing a significant increase in accessibility (adjusted p value < 0.05) were classified according to their log fold change (LogFC) in terms of chromatin accessibility in BECs exposed to flagellin (F) or not exposed (Ctl). These regions and their functionally linked genes were simultaneously summarised using the accessibility LogFC and expression fold changes (FC) from BECs exposed to F or not exposed (Ctrl). For each methodological approach, the average epigenomic and transcriptomic profiles (accessibility LogFC and expression FC, respectively) are shown per cluster. The values inside the boxes indicate the difference in the mean accessibility between Ctl and F at D2 and D6, estimated via analysis of variance per cluster. **C.** Schematic representation of average chromatin accessibility and gene expression changes for the three regulatory profiles defined using the clusters illustrated in 2B. Grey: Ctl condition, Red: F condition.

We then explored the effect of the increase in accessibility on the expression of neighbouring genes by associating these regions to their 15560 target genes according to the T-gene approach (18). The expression profiles were very similar for all clusters except for cluster 10 (**Figure 2A**) and coincide with the global expression changes previously shown (**Additional file 1: Fig. S1D**): a general upregulation during the time course for both conditions, with an enhanced and eventually a larger increase at D6F. However, this result does not indicate a particular correlation between gene expression and the epigenetic memory identified for some genomic regions (i.e., clusters 1, 2, 3, 5, 6, 8).

### Joint epigenomic and transcriptomic analyses discriminate sustained accessibility linked with expression changes upon flagellin pre-stimulation

To test the effect of epigenetic memory on the transcriptome, we jointly analysed accessibility and expression changes. Thus, we calculated the SOM for the complete dataset, including the regions with significant changes in accessibility and the expression of their target genes.

For the regions gaining accessibility, this implied the analysis of a high dimensional dataset, including 12173 links between 11877 regulatory regions and their 6094 potential target genes, under 5 conditions: D0 and two successive time points (D2 and D6) with or without pre-stimulation by flagellin. We again identified ten clusters on the resulting SOM that present, except for cluster 10, a sustained gain of accessibility that is stronger in F than in Ctl (**Figure 2B**).

Regions are now classified in a completely different way than in the previous unimodal analysis, with only accessibility information used as input data (**Additional file 1: Fig. S2A).** Additionally, loci gaining accessibility are associated with distinct gene expression profiles that allow for the identification of three regulatory profiles, as shown in **Figure 2C**: i) Memory profile corresponding to the upregulation of 636 genes either by F pre-stimulation alone (cluster 3) or with significantly greater expression upon stimulation (clusters 5 and 8); ii) Enhancement profile corresponding to 2794 genes upregulated but with no significant difference in expression between F and Ctl (clusters 1, 4, and 7); and iii) Downregulated profile corresponding to 2856 genes with similar (clusters 2 and 6) or different (clusters 9 and 10) expression changes between the Ctl and F conditions.

We performed the same analysis for the 10768 regions showing significant loss of accessibility under Ctl or F conditions and for their target genes (**Additional file 1: Fig. S2B**). Interestingly, the significance of the accessibility loss for F in comparison to Ctl is greater at D2 than at D6 for all clusters. This suggests that the initial changes are not sustained throughout the time course, and thus, there is no evidence for epigenetic memory upon stimulation for regions losing accessibility. In summary, the global changes induced by flagellin exposure that are maintained upon cell division mostly involve accessibility gain and not loss. Additionally, for some specific regions, a gain in chromatin accessibility implies a sustained increase in target gene expression, which is defined as the regulatory Memory profile.

### Genes clustered in the Memory profile specifically regulate the inflammatory response

To functionally characterise the three profiles defined above, we performed an enrichment analysis of the gene ontology (GO) biological processes for the 636, 2794, and 2856 target genes of the Memory, Enhancement and Downregulated profiles, respectively.

Enriched GO terms were chosen by setting a false discovery rate (FDR) < 0.05 and were sorted by the proportion of genes per profile annotated to a given term (**Supplementary Table 1**). We obtained 8, 25 and 18 enriched terms for the Memory, Enhancement and Downregulated profiles, respectively. Few terms were common to the 3 profiles. The terms “Defence response to virus” were shared by the Memory and Enhancement profiles, and “Cell migration” and “Protein phosphorylation” were shared by the Enhancement and Downregulated profiles. However, most GO terms were classified specifically into one regulatory profile. Indeed, 88%, 88% and 89% of the terms were regulated specifically in the Memory, Enhancement and Downregulated profiles, respectively.

Among the seven terms specific to the Memory profile, the highest proportion of target genes were classified as “Apoptotic process” (**Figure 3A)**. We validated this observation by PCR and confirmed that preexposure of BECs to flagellin significantly increased the expression of the DNA damage regulated autophagy modulator 1 (*DRAM1*), a gene involved in apoptosis, at D6F compared to D6 (5.3 ± 1 a.u. *vs.* 2.5 ± 0.6 a.u.) (**Figure 3B**). Interestingly, we found that five of the seven terms in the Memory profile were related to inflammation: “Innate immune”, “Inflammatory response”, “Cellular response to lipopolysaccharide”, “Positive regulation of inflammatory response”, and “Neutrophil chemotaxis” (annotated with * in Figure 3A). To validate these observations, we measured the expression of *CXCL6*, a gene annotated by three terms linked to inflammation namely, “Neutrophil chemotaxis,” “Cellular response to lipopolysaccharide,” and “Inflammatory response.” We observed a significant increase in *CXCL6* expression induced by flagellin preexposure at D2F *vs.* D2 (31.6 ± 7 a.u. *vs.* 4.17 ± 0.8 a.u.) that persisted at D6F *vs.* D6 (103.9 ± 34 a.u. *vs.* 17.0 ± 7.7 a.u.) (**Figure 3C**).

**Figure 3:**
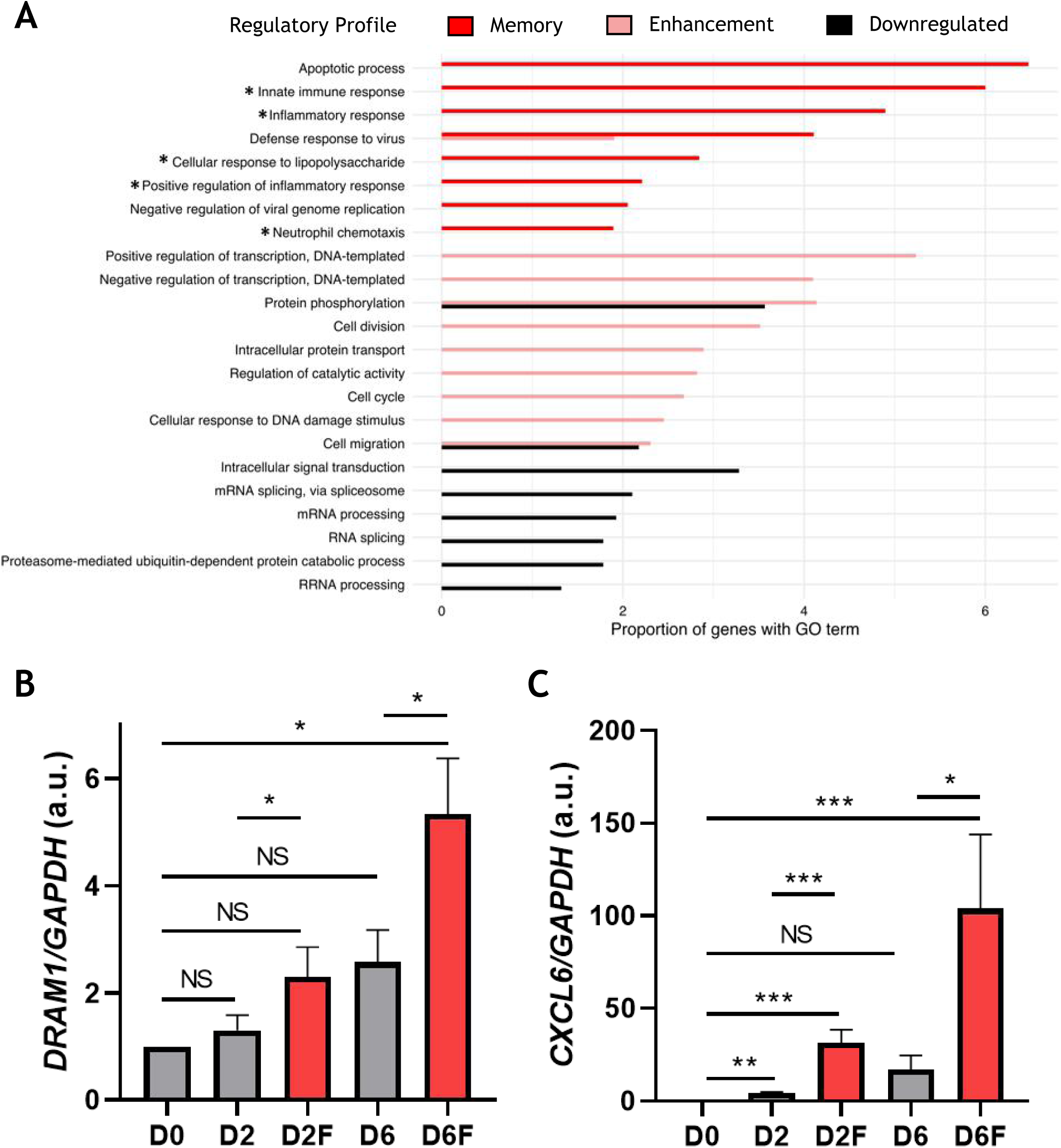
Functional analysis of the three regulatory profiles. **A.** Gene Ontology enrichment analysis was performed for genes belonging to the three regulatory profiles (Memory, Enhancement and Downregulated). The top enriched (FDR < 0.05) biological processes were selected and represented using the proportion of genes per profile annotated to a given GO term. **B, C.** *DRAM1* (**B**) and *CXCL6* (**C**) expression (in arbitrary units, a.u.) at days 0, 2 and 6 in BECs exposed to flagellin (F) or not exposed (Ctl). Each histogram represents the mean expression +/− standard error of the mean (n=5, ANOVA with pairwise comparison and Bonferroni’s multiple-comparison test, NS: not significant; *p<0.05; **p<0.01; ***p<0.001; ****p<0.0001).

Enriched terms specifically regulated in the Enhancement profile were mostly related to the basic metabolism of cells, such as “Regulation of transcription”, “Cell division”, “Cell cycle” and “Cellular response to DNA damage stimulus”. Terms enriched for the Downregulated profile point at functions involved in transcriptional and protein synthesis regulation, such as “mRNA splicing via spliceosome”, “mRNA processing”, “RNA splicing”, “Proteasome-mediated ubiquitin-dependent protein catabolic process”, and “rRNA processing”.

In conclusion, we showed that the three regulatory profiles defined via the joint analysis included sets of genes involved in distinct biological processes. Notably, genes in the Memory profile, which show upregulated expression and are associated with a persistent gain of accessibility upon flagellin pre-stimulation, are mostly involved in inflammation and innate immune processes.

### Epigenetic memory is associated with distinct regulatory networks

Transcription factors (TFs) are effectors of cell phenotype and function. As they recruit chromatin-modifying enzymes that modulate gene expression, they are cornerstones of epigenetic signalling (19). We thus aimed to identify the active transcription regulators to reconstruct the gene regulatory network involved in the induction of the training state. A footprinting analysis was performed for each regulatory profile using the HINT-ATAC method (20), where footprints denote protected regions within open chromatin peaks. We focused on TFs that exhibited changes in activity between the F and Ctl states throughout the time course. As the greatest changes in accessibility occurred towards the end of the time course, we specifically concentrated on changes between D2 and D6 (**Additional file 1: Fig. S1B**).

Hence, we focused on the predicted TF activity score by comparing the accessibility of their target sequence in F and Ctl in the three regulatory profiles. We found that the activity scores of the TFs differed among the three profiles and between clusters of the same profile (**Figure 4A**). We then focused on the TFs significantly activated by flagellin, which had the highest activity score (i.e., > 0.5) and were predicted to regulate genes in the Memory profile. We identified two major TF families: the heterodimeric complex activator protein-1 (AP-1) with JUN::JUNB, FOSL2::JUN, FOS::JUNB, and FOS::JUN in cluster 3; JUN::JUNB, FOS::JUNB, and FOSB::JUNB in cluster 5; FOS::JUN, FOSL2::JUN, and FOSL2 in cluster 8; and the interferon regulatory factor (IRF) family with IRF3 and IRF1 in clusters 5 and 8, respectively. EHF, DBP, Stat2, NFIB, GABPA and IKZF1 were also found in cluster 3; Foxd3, IRF3, IKZF1, and ELF3 were found in cluster 5; and EHF, NFIB, PRDM1, IRF1, SPIB, SP3, KLF9, JDP2, ETV4, EWSR1-FLI1 and ZNF384 were found in cluster 8. Among the genes predicted to be targeted by all TFs and belonging to the Memory profile, *CXCL1*, *CXCL6*, *NFKBIA*, *NAMPT* and *SLPI* were identified in cluster 3; *IL1B*, *DRAM1*, *CTSO*, *CLDN1* and *SERPINB3* were identified in cluster 5; and *CCL2*, *GBP2*, *RARRES1* and *TNFSF10* were identified in cluster 8. The relationships between TFs whose activity increased and that of the genes in the Memory profile are illustrated in Figure **4B** and are grouped in **Supplementary Table 2**.

**Figure 4:**
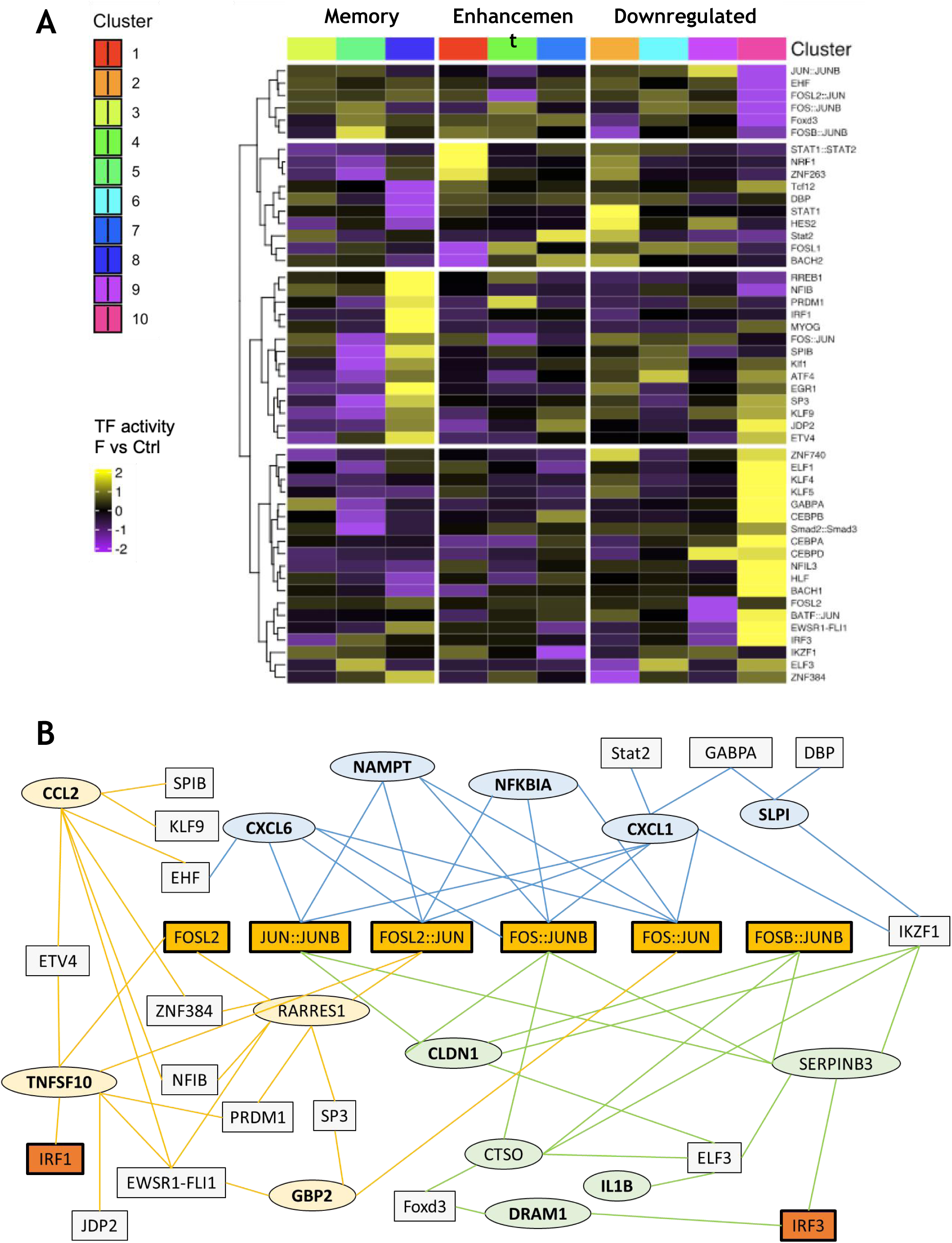
Transcription factor (TF) activity comparison between F and Ctl conditions per regulatory profile. **A.** TF activity was estimated from the ATACseq data using the footprinting approach implemented in HINT-ATAC. TF activity was analysed in BECs exposed to flagellin (F) and in BECs not exposed to flagellin (Ctl) by comparing their activity between days 6 and 2 per condition. For each cluster, the top 10 TFs with the greatest expression difference between F and Ctl and a number of binding sites in the highest 5% quantile were selected. **B.** Schematic representation of the predicted regulatory network of Memory clusters. The predicted TFs (squares) and their target genes (circles) detected in the transcriptomic data are represented by links, as one gene can be targeted by several TFs and one TF can target several genes. Genes with a light blue background belong to cluster 3, genes with a light orange background belong to cluster 8, genes with a light green background belong to cluster 5, and finally, the genes written in bold correspond to those involved in innate immunity.

## Discussion

It is now well established that innate immune memory occurs through epigenetic and metabolic reprogramming (12,13,21,22). Given our previous results that suggest a link between immune memory in BECs and epigenetic mechanisms, particularly histone modifications (12), we wanted to analyse changes in chromatin condensation induced by flagellin priming that sustain the reprogramming of transcriptional patterns. We compared transcriptomic and epigenomic changes following flagellin priming over a 6-day time course. This led to the analysis of a high-dimensional dataset, including ∼12K links between thousands of regulatory regions and their potential target genes, under 5 conditions, at D0 and two successive time points (D2 and D6) with or without flagellin pre-stimulation. A major lesson learned using our methodological approach is the relevance of the joint analysis of the epigenome and transcriptome to obtain a classification of dynamic regulatory regions that are associated with expression changes and can be interpreted in light of the biological hypothesis being tested. We applied an unsupervised approach, self-organising maps (SOMs), to identify regulatory modules, i.e., regions and linked genes, with different behaviours between the unstimulated and prestimulated time courses. This method produces a low-dimensional representation that summarises the linear and nonlinear relationships among the initial high-dimensional data points.

With respect to the unimodal approach, we focused on the increase in chromatin accessibility. SOM analysis revealed profiles with a similar trend, i.e., a more important increase in accessibility for the stimulated time course with a range of accessibility changes per profile, varying from 0 to almost 4 log fold. However, this variability is not mirrored by distinct changes in the expression of the linked genes, which on average tended to be more highly expressed in the prestimulated time course without significant differences among profiles. This result coincides with the well-known correlation between accessible chromatin and actively transcribed genes and has no connection with the establishment of innate memory.

The SOM outcome was completely changed following a joint analysis of accessibility and expression, allowing the transcriptome to guide the classification of changes over two time courses. This allowed for the identification of distinct expression profiles with accessibility changes bounded between 0 and 2 log fold. Such expression profiles have an identity that goes beyond their quantitative similarity: they have three qualitatively distinct behaviours, i.e., Memory, Enhancement and Downregulation, which, moreover, are enriched in different biological functions. These results suggest that flagellin induces a general nonspecific decondensation of chromatin. Around some specific genes, such modulation induces epigenomic changes that subsist across multiple cell divisions. They are not necessarily statistically significant at D2 but increase steadily from D0 to D6. Interestingly, genes related to this memory-like behaviour are mostly involved in inflammatory and innate immune responses. This finding was further validated by the overexpression of two genes, DRAM1 and CXCL6, included in the Memory profile and associated with open chromatin clusters. The hypothesis that flagellin exposure plays a role in establishing innate immune memory was reinforced by the absence of a flagellin-specific effect on the expression of genes in the Enhancement profile. Indeed, genes clustered in the Enhancement profile are involved in basic cell metabolism, and the upregulation of their expression was independent of flagellin stimulation. Genes clustered in the Downregulated profile had downregulated expression even though they were associated with gained loci.

Epithelial cells are continuously subject to regeneration due to permanent cell division. In our experimental procedure, only 30% of the BECs analysed at D6 (10.3 ×10^4^ cells/well) were in contact with flagellin at D0 (2.8 ×10^4^ cells/well). Therefore, ATAC-seq analysis was performed with respect to cell concentration at D0, D2 and D6 to avoid bias related to the increasing number of cells. Moreover, we found that flagellin exposure induced epigenetic modifications at D2 that persisted until D6, even though the cells were washed in between D2 and D6, and the stimulus was thus removed. Therefore, the memory phenotype is retained over time across cell divisions, suggesting that mechanisms involved in BEC memory are transmitted to daughter cells over generations. Interestingly, Sun and his colleagues also showed that macrophages were still transcriptionally, epigenetically, and functionally reprogrammed after 14 rounds of cell division following stimulus washout. Moreover, they demonstrated that long-lasting epigenetic modifications in trained cells were always coupled with changes in transcription factor (TF) activity, suggesting that TF plays a central role in driving the transmission of epigenetic changes across cell divisions (23). By exploring the relationships between TFs and durable epigenetic programs in the Memory profile, we predicted several TFs associated with open chromatin that correlate with upregulated gene expression. TF analysis suggested that the training state relies on the remodelling of TF activity since coordinated changes affect multiple members of many TF families. Differential TF motif accessibility analysis across the time course in prestimulated and unprestimulated BECs revealed a central role for the TFs Fos/Jun (AP-1) and IRF family motifs in the expression of genes included in the Memory profile. The IRF family consists of nine members in humans, IRF1-9, which play central roles in the production of interferons. AP-1 is a heterodimeric TF comprising proteins belonging to the JUN, FOS, ATF, MAF and CREB families. AP-1, which controls both basal and inducible transcription of several genes, is involved in the pathways activated upon binding of flagellin to its extracellular receptor, TLR5 (24). The AP-1 and IRF families are involved in trained immunity. Indeed, an investigation of epigenetic innate immune memory resulting from SARS-CoV-2 infection performed by analysing chromatin and transcription in monocytes and their progenitors revealed an increase in the activity of AP-1 and IRF (25). Recent studies have also revealed a key role for AP-1 in chromatin accessibility. Therefore, Vierbuchen *et al*. provided evidence that AP-1 TFs, together with other specific TFs, bind to nucleosome-occluded enhancers via the recruitment of the BAF complex, which induces nucleosome remodelling, and establish an accessible chromatin state (26). In a murine model of trained immunity developed by chronic skin inflammation, Larsen *et al.* showed that cells exposed to inflammatory mediators are enriched in STAT3 and FOS-JUN at memory domains, defined as chromatin regions that gain accessibility during the inflammatory response and remain so following resolution (27).

Our results provide evidence that flagellin exposure-induced innate immune memory in BECs is the result of chromatin decondensation of a subset of specific regions. The identification of such regions and the putative drivers of this process was possible thanks to our methodological approach. The joint unsupervised analysis of the epigenome and transcriptome allowed us to identify regulatory modules and highlighted a Memory profile specifically induced after flagellin exposure. Under these conditions, open chromatin was associated with the upregulation in expression of genes involved mainly in apoptosis, inflammation and the response to infection pathways. Although our data did not allow us to delineate the specific timing of these fine-tuned series of events and which events are specifically required for memory transmission in dividing cells, they reveal a multitude of potential targets for modulating the response of BECs in the context of innate immune memory.

## Conclusion

In conclusion, we demonstrated that BECs can be epigenetically reprogrammed in a sustained way after a primary microbial contact, a phenomenon that modulates their response to further infection. The epigenetic mechanisms that sustain this phenomenon should be further analysed to identify the chromatin modifiers that contribute to regulating signalling pathways and altering transcription. This step is mandatory for potential immune control in the management of pulmonary pathologies. Indeed, suppressing or preventing trained immunity could be a relevant option in the treatment of chronic pulmonary diseases associated with a chronic inflammatory status, such as cystic fibrosis characterised by chronic pulmonary infections and inflammatory exacerbations. Conversely, the development of trained immunity inducers could be useful for the treatment of immunosuppressed patients, with chemotherapy treatments, for example, to prevent the development of pulmonary infections. Management of memory duration by epigenetic modulators is another avenue to be investigated. All these future approaches will aid in the development of innovative therapeutics.

## Methods

### Cell preparation and stimulation

Human BECs (BEAS-2B cell line; American Type Culture Collection, Rockville, MD, USA) were maintained at 37 °C and 5% CO2 in F12 medium (Invitrogen, Waltham, MA, USA) supplemented with 10% foetal calf serum, 1% penicillin‒streptomycin, and 10 mM HEPES and stimulated in F12 without antibiotics. BECs were cultured in 96-well plates at 1.5 × 10^4^ cells per well on day −1. On day 0, BECs were stimulated with or without ultrapure *P. aeruginosa* flagellin (InvivoGen, Waltham, MA, USA) (5 μg/mL) for 48 h. On day 2, the cells were washed twice and incubated with medium for a resting period of 4 days until day 6. For DNA and RNA extraction, BECs were lysed at D0 just before pre-stimulation with flagellin, at D2 at the time of washing, and at D6 after the resting period (**Figure 1**). All experiments were performed in triplicate. The DNA preparation protocol used a concentration extracted from 5 × 10^4^ cells. Because the BECs multiplied during the time course of the memory experiment, the cells were counted at D0 (2.8 ×10^4^ cells/well), D2 (4.5 ×10^4^ cells/well) and D6 (10.3 ×10^4^ cells/well). Therefore, at D0 and D2, DNA was extracted from 5 and 2 pooled wells of cultured BECs, respectively, and BECs from only one well were used for DNA extraction at D6. For RNA extraction, we performed the same procedure to obtain approximately 3 µg at 100 ng/µL at each time point. Therefore, 30, 12 and 10 wells were pooled to extract RNA at D0, D2 and D6, respectively.

To ensure that BECs were in a training state after flagellin exposure, we performed an experiment in which BECs were pre-stimulated with or without flagellin from D0 to D2 and stimulated on D6 with live *A. fumigatus* conidia at a multiplicity of infection (MOI) of 2 for 15 h. The *A. fumigatus* DAL strain (CBS 144.89) was grown as described elsewhere (28). Following *A. fumigatus* infection, the culture supernatant was recovered, and IL-8 and IL-6 levels were quantified with an enzyme-linked immunosorbent assay (R&D Systems, Minneapolis, MN, USA).

### RNA purification and DNA transposition

RNA was isolated using a NucleoSpin RNA/Protein kit (Macherey Nagel, Duren, Germany). For the preparation of the DNA, we used a protocol inspired by Buenrostro *et al*. (29). Briefly, adherent BECs were lysed with 20 µL/well ATAC-resuspension buffer (RSB) (10 mM Tris-HCl pH 7.4, 10 mM NaCl, 3 mM MgCl2) supplemented with 0.1% NP40 (Sigma Aldrich), 0.1% Tween-20 (Sigma Aldrich) and 0.01% digitonin (Promega), incubated for 3 min on ice, and washed with 0.1 mL of cold ATAC-RSB supplemented with 0.1% Tween-20 without NP-40 or digitonin. The extracted nuclei were pelleted at 500 × g for 10 min at 4 °C. The pellets were resuspended in 50 µL of transposition mixture (20 mM Tris-HCl pH 7.6, 10 mM MgCl2, 20% dimethyl formamide, 100 nM transposase, 1% digitonin, 0.1% Tween-20) and incubated at 37 °C for 30 min in a thermomixer with mixing at 1000 RPM. Then, the fragmented DNA was purified using a DNA Clean and Concentrator-5 Kit (Zymo Research, #ZYM-D4014, Irvine, CA, USA) eluted with 21 µL of elution buffer and stored at −20 °C.

### Library generation and sequencing

DNA libraries for ATAC-Seq were prepared at the Institute of Cochin’s GENOM’IC facility from transposed DNA using the Nextera® DNA kit from Illumina, following the final amplification by PCR step instructions. The reaction tubes contained 15 µL Nextera PCR Master Mix, 5 µL Illumina PCR Primer Cocktail, 5 µL Index Primer i7, 5 µL Index Primer i5, 10 µL H_2_O and 10 µL transposed DNA. PCR was performed with the following parameters: 3 min at 72 °C, 30 s at 98 °C, and eight cycles of 10 s at 98 °C, 30 s at 60 °C, and 2 min at 72 °C. PCR products were purified using AMPureXP™ beads from Agencourt at a 1X ratio. The final library quality was assessed on a DNA High Sensitivity™ chip on a Bioanalyzer 2100 from Agilent, and the concentration was measured using a Qubit dsDNA HS Assay™ kit from Thermo Fisher Scientific (the average yield was 12.1 ng/µL). The molarity of each library was calculated with the following formula: Qubit measure * 1000000/((average peak size in bp * 607.4) + 157.9).

Each library was diluted to a concentration of 10 nM, and a final pool was set at 2 nM prior to sequencing.

Sequencing of the ATAC-Seq libraries was performed on a NextSeq™ 500 from Illumina (GENOM’IC). The sequencer parameters were set as follows: read 1 = 76 cycles, index 1 = 8 cycles, index 2 = 8 cycles, and read 2 = 76 cycles (using a NextSeq 500 High Output 150 cycles reagent kit from Illumina). The internal PhiX control was added to the library pool to reach 1% of the total reads. The sequenced pool was diluted to a 2 pM concentration and loaded in the sequencer.

The 15 libraries were successfully sequenced. The total number of reads after quality filtering from the sequencer reached 419,765,351, with a Q30 of 93.1% and a cluster density of 192.57 clusters/mm². PhiX reads accounted for 1.86% of the total, as expected. The number of reads per sample averaged 25.34 M passing filter reads. Demultiplexing and quality of the sequences were performed using the Aozan tool from ENS, Paris (30), and all quality controls were satisfactory. Resequencing was needed for eight samples. Sequencing was performed as previously described and yielded 527,141,088 passing filter reads with a Q30 of 91.72% and a cluster density of 240.03 clusters/mm². PhiX control reached 1.05% of the total reads. The number of reads per sample averaged 61.10 M passing filter reads. Demultiplexing and quality of the sequences were performed using the Aozan tool as previously described, and the results were satisfactory.

### Microarray and differential expression analysis

RNA was isolated using a NucleoSpin RNA/Protein kit (Macherey Nagel, Duren, Germany). One hundred ng total RNA was reverse transcribed following the GeneChip® WT Plus Reagent Kit (Affymetrix). Briefly, the resulting double-stranded cDNA was used for *in vitro* transcription with T7 RNA polymerase (all these steps were included in the WT cDNA synthesis and amplification kit of Affymetrix). After purification according to the Affymetrix protocol, 5.5 µg of Sens Target DNA were fragmented and biotinylated. After control of fragmentation using Bioanalyzer 2100, cDNA was hybridised to a GeneChip® ClariomS Human (Life Technologies, Carlsbad, Californie, USA) at 45 °C for 17 h. After overnight hybridisation, the chips were washed on a FS450 fluidic station following specific protocols (Affymetrix) and scanned using a GCS3000 7G.

The scanned images were then analysed with Expression Console software (Affymetrix) to obtain the raw data (CEL files) and quality control metrics. The observations of some of these metrics and the study of the distribution of the raw data showed no outlier data. RMA normalisation was performed using R with version 23 of the Entrezgene CDF brain array.

Changes in gene expression between conditions A and B were measured using a fold change (FC) defined as follows:

expression ratio = expression in A/expression in B
FC = expression ratio if expression ratio > 1
FC = −1/expression ratio if 0 < expression ratio < 1.

### ATAC-seq analysis

The data were analysed with ePeak (31). Reads were mapped against the human genome (hg38 from Ensembl release 94) using Bowtie2 v2.1.0 (32) with custom parameters (-X2000 --dovetail --no-mixed --no-discordant --end-to-end --very-sensitive). Duplicated reads were removed with markDuplicates from Picard tools v1.94, and blacklisted regions were excluded using bedTools intersectBed v2.17. Nucleosome-free fragments (fragments smaller than 120 base pairs) were selected using the awk command. Accessible peaks were called using MACS2 v2.1.0 (33) with the following parameters: --nomodel --extsize=100 -p 0.1. Reproducible peaks were selected using the irreproducible discovery rate (IDR) method as described by (34). A union peak file was created by merging the reproducible peak list for all the conditions of the experiment using bedTools merge with default parameters. A count matrix was generated using this union peak file and all BAM files with featureCounts v1.4.6-p3 (35) from the Subreads package.

### Differential peak analysis

The count matrix was analysed using R version 4.0.2 and the Bioconductor package DESeq2 version 1.28.0 (36). The normalisation and dispersion estimation were performed with DESeq2 using the default parameters, and statistical tests for differential expression were performed using the independent filtering algorithm. A generalised linear model was used to test for differential accessibility between D0, D2 and D6, with or without pre-stimulation by flagellin. For each pairwise comparison, the raw *P* values were adjusted for multiple testing according to the Benjamini and Hochberg (BH) procedure (37), and peaks with an adjusted *P* value less than 0.05 were considered differentially accessible.

### Summary and classification of accessibility and transcriptomic data

Accessibility (from ATAC-seq) logFold changes and transcriptomic (from Microarray) fold changes were calculated between D2 and D0 and between D6 and D0 for the control and stimulated conditions. Peaks gaining accessibility (fold change between D6F and D0 > 0) or losing accessibility (fold change between D6F and D0 < 0) were analysed separately to facilitate the biological interpretability of the results. We then predicted the regulatory links between differentially accessible peaks and genes using the T gene and selected the most significant links (distance < 500 kb and correlation p value < 0.05). Finally, we used self-organising maps (SOM), an unsupervised machine learning technique (38), to obtain a low-dimensional representation of our high-dimensional dataset, including peaks and genes, over the 4 comparisons described above. We used the SOM implementation of the Kononen R package version 3.0.12 with a 24*24 hexagonal topological grid. On the obtained SOM, we performed a k-means classification to obtain regulatory profiles. Clustering was stopped at 10 clusters to ensure a balance between granularity and biological interpretability. SOMs were calculated using two approaches: only on differentially accessible peaks (unimodal analysis) and the use of both differentially accessible peaks and their linked genes, as explained above (joint analysis).

For each regulatory profile (accessibility gain and loss), we plotted the accessibility logFold changes and the expression fold changes per cluster. *P* values between stimulated and control conditions were computed by fitting an analysis of variance model of the logFC values and the Ctl or F condition.

### Gene function annotation and gene enrichment analysis

To identify the biological functions characterising each regulatory profile, we performed an enrichment analysis of Gene Ontology (GO) biological process terms using the DAVID knowledgebase (39). Enriched GO terms were selected by setting a false discovery rate (FDR) < 0.05 and were sorted by the proportion of genes annotated to each term.

### Differential transcription factor activity

Differential transcription factor (TF) footprinting was performed using HINT-Atac v0.13.1 (20) as described here: https://reg-gen.readthedocs.io/en/latest/hint/tutorial-dendritic-cell.html. To increase the number of paired reads per sample to ∼200M, we merged all replicates per time point and condition (Ctl and F). The regions assigned to each TF were used as inputs for the footprint analysis. Footprints were called with ‘rgt-hint footprinting’ with the following parameters: --atac-seq --paired-end. Then, TF binding motifs were searched among the predicted footprints using the ‘rgt-motifanalysis matching’ command. Finally, the average ATAC-seq coverage was computed for each TF and compared to that under other conditions with the ‘rgt-hint differential’ command. For each TF, an activity score was calculated as described in Li *et al*. 2019 (20) by adding the tag counts and protection score for each condition. The activity scores of F and Ctl were compared by dividing the activity score difference for F (D6F-D2F) and Ctl (D6-D2). For each regulatory profile, the top 10 TFs with the largest differences between F and Ctl activity and the greatest number of binding sites in the highest 5% quantile were selected. Heatmaps representing differential activity scores between F and Ctl and for each cluster were plotted using the ComplexHeatmap package version 2.16 (40).

### RT-qPCR

RNA was isolated using a NucleoSpin RNA/Protein kit (Macherey Nagel, Duren, Germany). RT was performed using a high-capacity cDNA kit (Applied Biosystems, Foster City, CA, USA). Real-time qPCR was performed with an ABI StepOnePlus using TaqMan Fast Universal PCR Master Mix (Applied Biosystems), TaqMan probes for *GAPDH* (Hs_2786624), *CXCL6* (Hs_06632196), and *DRAM1* (Hs_01022842), and cDNA as a template. For relative quantification, the amount of target genes was normalized to the expression of *GAPDH* relative to that of the control cells used as a calibrator and was calculated using the 2^−ΔΔCt^ method.

## Abbreviations

ATAC-Seq: Assay for Transposase-Accessible Chromatin with High-throughput Sequencing
BECs: Bronchial epithelial cells
Ctl: Control
D: Day
F: flagellin
FC: Fold change
FDR: false discovery rate
NES: normalized enrichment score
SOMs: self-organizing maps
TF: Transcription factor

## Declarations

### Ethics approval and consent to participate

Not applicable

### Consent for publication

Not applicable

### Availability of data and materials

The data discussed in this publication have been deposited in NCBI’s Gene Expression Omnibus (41) and are accessible through GEO Series accession number GSE227261 (https://www.ncbi.nlm.nih.gov/geo/query/acc.cgi?acc=GSE227261). The code used to generate the figures presented in this manuscript can be accessed via the following link: https://gitlab.pasteur.fr/rlegendr/bec_memory.

### Authors’ contributions

JB and VB conceived and performed the experiments. MLG and JYC helped to conceive the experimental protocol. JH and SJ performed the sequencing of the data and performed the differential analysis of the microarray. RL and CC performed the bioinformatics and biostatistics analyses. JB, RL, VB and CC analysed and interpreted the results. JB, RL, VB and CC wrote the manuscript. All the authors have read and approved the final manuscript.

## Acknowledgements

The authors would like to thank Vincent Guillemot for his statistical input and all members of the Hub of Bioinformatics and Biostatistics for useful discussions. This work used the computational and storage services (MAESTRO cluster) provided by the IT department at the Institut Pasteur, Paris.

## Supplementary Figure legends

**Additional file 1: Fig. S1: Differential chromatin accessibility and gene expression analysis. A.** Differentially accessible regions were calculated for the D2 *vs.* D0, D2F *vs.* D0, D6 *vs.* D0 and D6F *vs.* D0 comparisons and retained if the adjusted p value was lower than 0.05. The UpSet plot shows the intersections across all comparisons: horizontal bars indicate the total number of differentially accessible regions, and vertical bars indicate the number of regions that are unique to a given comparison or common between two or more comparisons, as indicated by the dots underneath. **B.** Representation of the 500 most variable accessible regions on the three first principal components. The colours indicate the different time points, and the percentages on each axis specify the percentage of variability explained by a given component. **C.** UpSet plot representing differentially expressed genes across comparisons as in A. **D.** Representation of the 500 most variably expressed genes as in B.

**Additional file 1: Fig. S2: Changes in the epigenome and transcriptome associated with regions losing chromatin accessibility during BEC priming. A.** Comparison of clustering results using only accessibility (left) or both accessibility and expression data (right). Stacked bar plots indicate the distribution of regions among clusters, while flow streams highlight their differential classification between the two methodological approaches. **B.** Regions exhibiting significant loss of accessibility (adjusted p value < 0.05) were classified using the log fold changes (logFC) in chromatin accessibility between BECs exposed to flagellin (F) and those not exposed to flagellin (Ctl). These regions and their functionally linked genes were simultaneously summarised using the accessibility LogFC and expression fold changes (FCs) from BECs exposed to F or Ctl. The average epigenomic and transcriptomic profiles (accessibility logFC and expression FC, respectively) are shown per cluster. The values inside the boxes indicate the difference in the mean accessibility between the Ctl and F timecourses at D2 and D6, estimated via analysis of variance per cluster.

**Supplementary Table 1: Tables of enriched terms for the Memory, Enhancement and Downregulated profiles.** Enriched GO terms were chosen by setting a fold discovery rate (FDR) < 0.05 and were sorted by the proportion of genes per profile annotated to a given term. Families whose names appear on a grey background correspond to families that are common to different regulatory profiles.

**Supplementary Table 2: Table of the predicted relationships between the identified transcription factors and genes of the Memory profile.** Genes were sorted by cluster. Transcription factors whose predicted activity was significantly elevated in the presence of flagellin were placed on the same line as the genes whose transcription they regulate.

