## Supplementary Figures for "The epigenomic landscape of bronchial epithelial cells reveals the establishment of trained immunity"

### Supplementary Figure 1

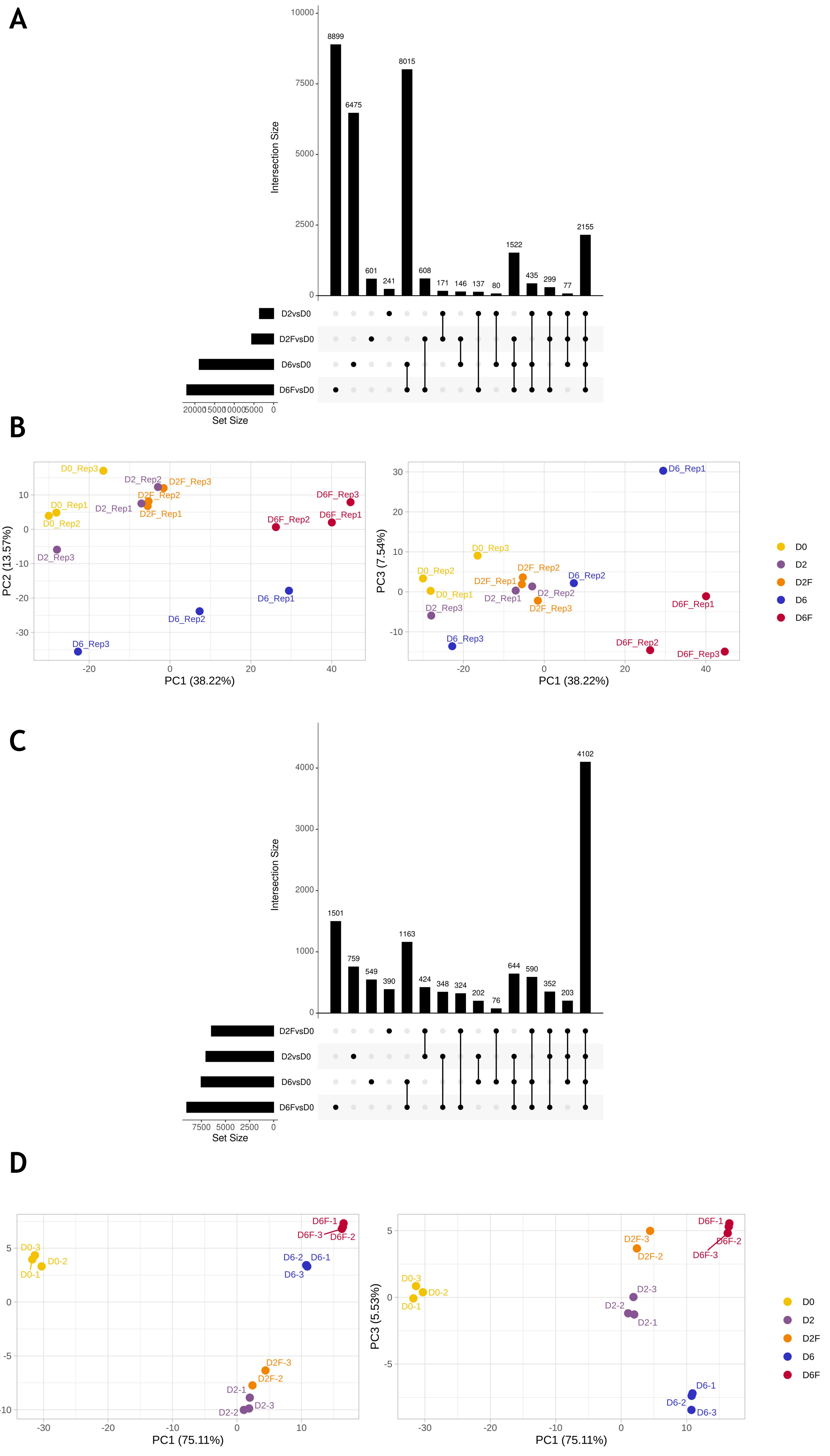

### Supplementary Figure 2

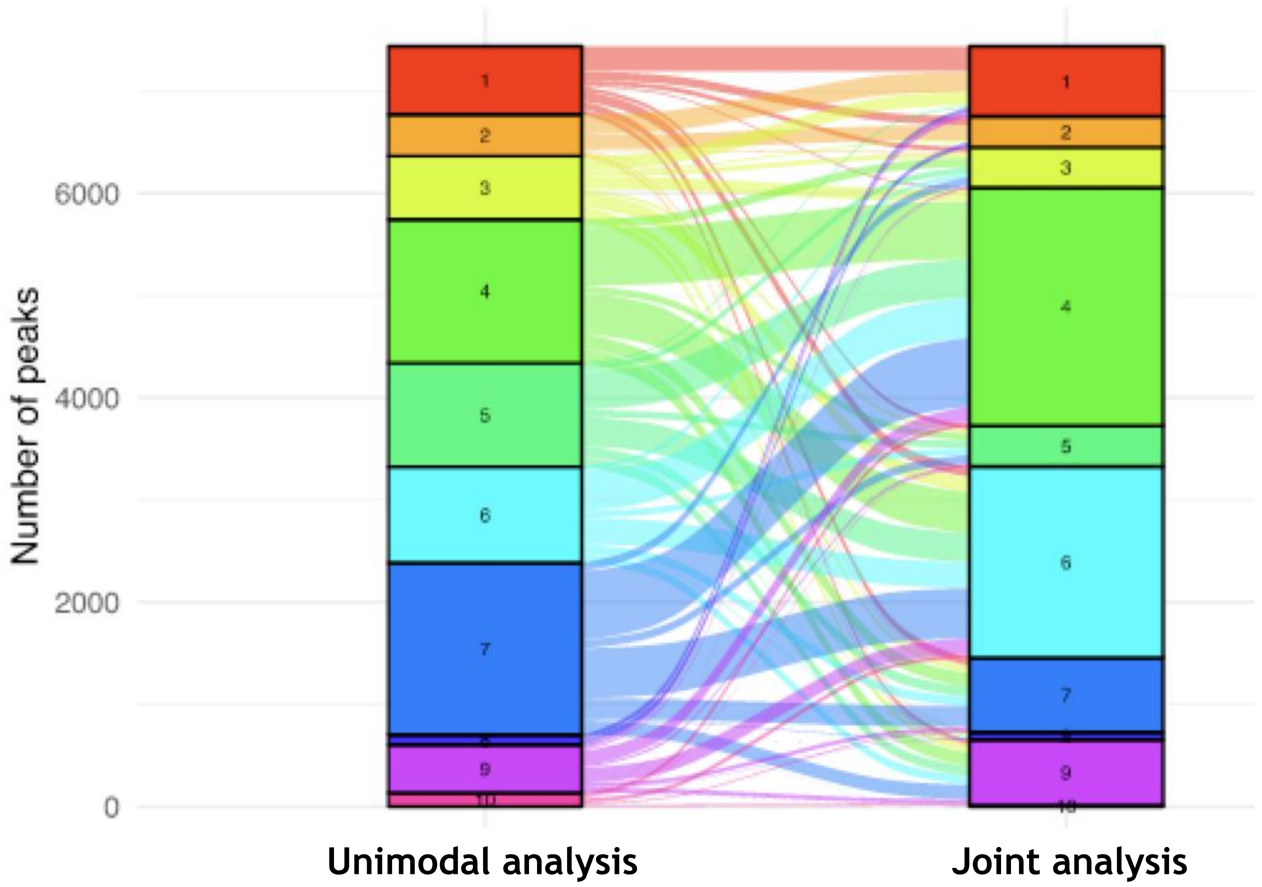


**B**

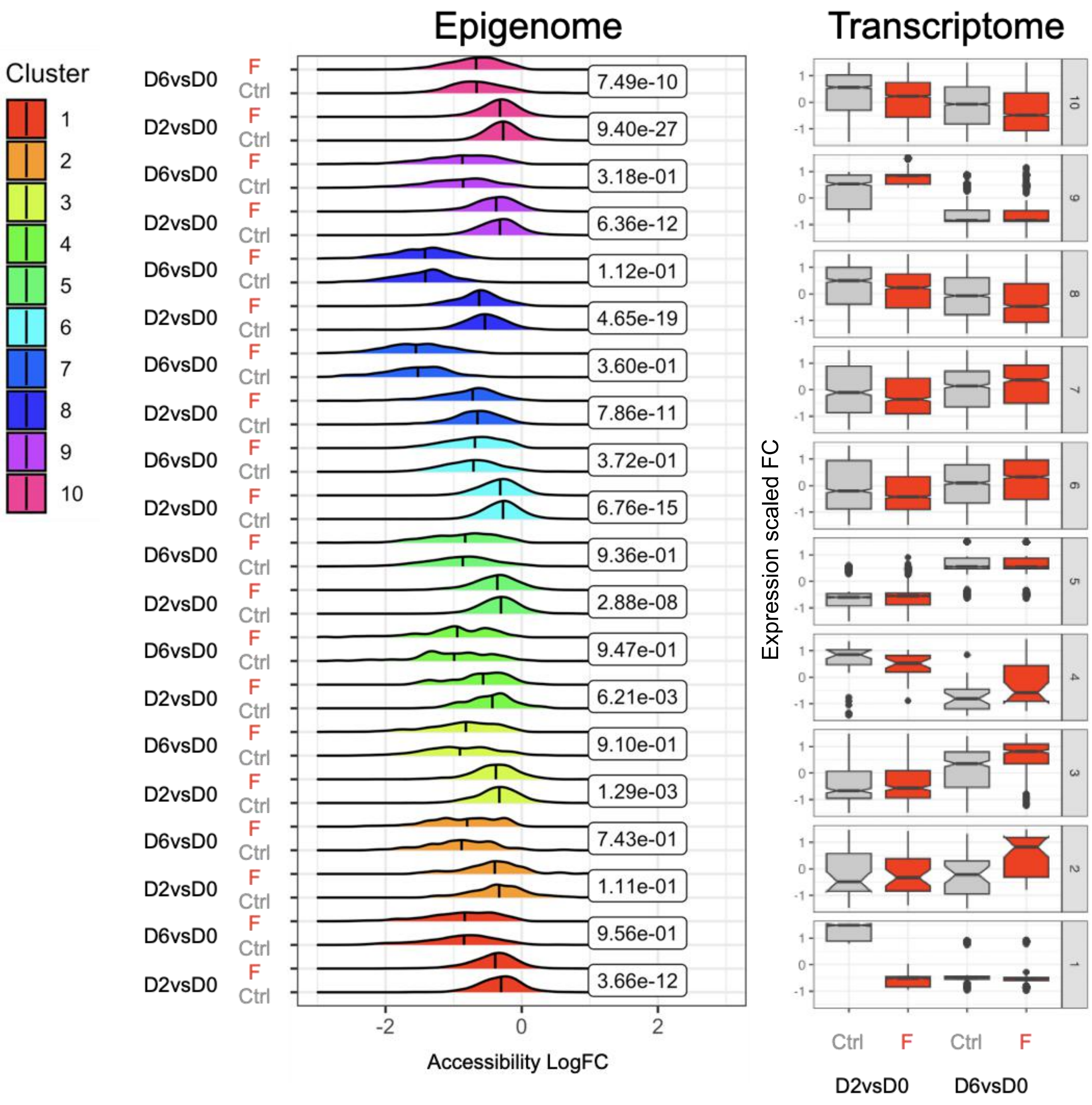

### Supplementary Table 1

| Memory | Biological Pathways | % | Fold Enrichment | FDR |
|---|---|---|---|---|
| GO:0006915 | apoptotic process | 6,48 | 2,17 | 0,0073 |
| GO:0045087 | innate immune response | 6,00 | 2,02 | 0,0338 |
| GO:0006954 | inflammatory response | 4,90 | 2,41 | 0,0127 |
| GO:0051607 | defense response to virus | 4,11 | 3,57 | 0,0001 |
| GO:0071222 | cellular response to lipopolysaccharide | 2,84 | 3,00 | 0,0451 |
| GO:0050729 | positive regulation of inflammatory response | 2,21 | 4,05 | 0,0235 |
| GO:0045071 | negative regulation of viral genome replication | 2,05 | 9,08 | 0,0000 |
| GO:0030593 | neutrophil chemotaxis | 1,90 | 4,70 | 0,0235 |

| Enhancement | Biological Pathways | % | Fold Enrichment | FDR |
|---|---|---|---|---|
| GO:0045944 | positive regulation of transcription from RNA polymerase II promoter | 8,42 | 1,46 | 0,0000 |
| GO:0000122 | negative regulation of transcription from RNA polymerase II promoter | 7,03 | 1,52 | 0,0000 |
| GO:0045893 | positive regulation of transcription, DNA-templated | 5,23 | 1,52 | 0,0002 |
| GO:0006468 | protein phosphorylation | 4,14 | 1,73 | 0,0000 |
| GO:0045892 | negative regulation of transcription, DNA-templated | 4,10 | 1,46 | 0,0149 |
| GO:0051301 | cell division | 3,51 | 1,93 | 0,0000 |
| GO:0006886 | intracellular protein transport | 2,89 | 1,84 | 0,0001 |
| GO:0050790 | regulation of catalytic activity | 2,82 | 1,57 | 0,0207 |
| GO:0007049 | cell cycle | 2,67 | 1,55 | 0,0371 |
| GO:0006974 | cellular response to DNA damage stimulus | 2,45 | 1,84 | 0,0004 |
| GO:0006281 | DNA repair | 2,34 | 1,67 | 0,0152 |
| GO:0016477 | cell migration | 2,31 | 1,79 | 0,0028 |
| GO:0043123 | positive regulation of I-kappaB kinase/NF-kappaB signaling | 1,98 | 2,16 | 0,0001 |
| GO:0018105 | peptidyl-serine phosphorylation | 1,90 | 2,19 | 0,0001 |
| GO:0051607 | defense response to virus | 1,90 | 1,71 | 0,0371 |
| GO:0006351 | transcription, DNA-templated | 1,87 | 1,77 | 0,0207 |
| GO:0006470 | protein dephosphorylation | 1,68 | 2,29 | 0,0001 |
| GO:0000278 | mitotic cell cycle | 1,65 | 2,24 | 0,0002 |
| GO:0046777 | protein autophosphorylation | 1,61 | 1,86 | 0,0208 |
| GO:0065003 | macromolecular complex assembly | 1,46 | 2,02 | 0,0093 |
| GO:0090630 | activation of GTPase activity | 1,21 | 1,98 | 0,0438 |
| GO:0006260 | DNA replication | 1,17 | 2,03 | 0,0371 |
| GO:0030509 | BMP signaling pathway | 0,95 | 2,27 | 0,0318 |
| GO:0010971 | positive regulation of G2/M transition of mitotic cell cycle | 0,51 | 3,71 | 0,0121 |
| GO:0043001 | Golgi to plasma membrane protein transport | 0,48 | 3,44 | 0,0371 |

| Downregulated | Biological Pathways | % | Fold Enrichment | FDR |
|---|---|---|---|---|
| GO:0006468 | protein phosphorylation | 3,57 | 1,47 | 0,0279 |
| GO:0035556 | intracellular signal transduction | 3,28 | 1,51 | 0,0274 |
| GO:0016477 | cell migration | 2,18 | 1,67 | 0,0279 |
| GO:0000398 | mRNA splicing, via spliceosome | 2,10 | 2,18 | 0,0001 |
| GO:0006397 | mRNA processing | 1,93 | 1,81 | 0,0120 |
| GO:0008380 | RNA splicing | 1,78 | 1,85 | 0,0120 |
| GO:0043161 | proteasome-mediated ubiquitin-dependent protein catabolic process | 1,78 | 1,75 | 0,0395 |
| GO:0006364 | rRNA processing | 1,32 | 2,10 | 0,0120 |
| GO:0032543 | mitochondrial translation | 1,18 | 2,55 | 0,0017 |
| GO:1903078 | positive regulation of protein localization to plasma membrane | 0,75 | 2,73 | 0,0179 |
| GO:0048146 | positive regulation of fibroblast proliferation | 0,71 | 2,56 | 0,0498 |
| GO:0006413 | translational initiation | 0,68 | 2,66 | 0,0466 |
| GO:0006446 | regulation of translational initiation | 0,57 | 3,83 | 0,0067 |
| GO:0050870 | positive regulation of T cell activation | 0,57 | 3,60 | 0,0106 |
| GO:0002503 | peptide antigen assembly with MHC class II protein complex | 0,39 | 5,10 | 0,0106 |
| GO:0002381 | immunoglobulin production involved in immunoglobulin mediated immune response | 0,39 | 4,80 | 0,0120 |
| GO:0002504 | antigen processing and presentation of peptide or polysaccharide antigen via MHC class II | 0,39 | 4,08 | 0,0412 |
| GO:0034383 | low-density lipoprotein particle clearance | 0,36 | 5,30 | 0,012[illegible] |

### Supplementary Table 2

| Clusters | Genes | Transcription factors | | | | | | | |
|---|---|---|---|---|---|---|---|---|---|
| 3 | CXCL1 | FOSL2::JUN | FOS::JUNB | FOS::JUN | JUN::JUNB | EHF | GABPA | IKZF1 | Stat2 |
| | CXCL6 | FOSL2::JUN | FOS::JUNB | FOS::JUN | JUN::JUNB | EHF | | | |
| | NFKBIA | FOSL2::JUN | FOS::JUNB | FOS::JUN | | | | | |
| | NAMPT | FOSL2::JUN | FOS::JUNB | FOS::JUN | JUN::JUNB | | | | |
| | SLPI | DBP | GABPA | IKZF1 | | | | | |
| 5 | IL1B | ELF3 | | | | | | | |
| | DRAM1 | Foxd3 | IRF3 | | | | | | |
| | CTSO | FOS::JUNB | FOSB::JUNB | Foxd3 | IKZF1 | ELF3 | | | |
| | CLDN1 | FOS::JUNB | FOSB::JUNB | JUN::JUNB | IKZF2 | ELF4 | | | |
| | SERPINB3 | FOS::JUNB | FOSB::JUNB | JUN::JUNB | IKZF3 | ELF5 | IRF3 | | |
| 8 | CCL2 | EWSR1-FLI1 | NFIB | SPIB | KLF9 | ETV4 | EHF | ZNF384 | |
| | GBP2 | EWSR1-FLI2 | FOS::JUN | SP3 | | | | | |
| | RARRES1 | EWSR1-FLI3 | FOSL2::JUN | FOSL2 | NFIB | PRDM1 | SP3 | ZNF384 | |
| | TNFSF10 | EWSR1-FLI4 | FOSL2::JUN | FOSL3 | PRDM1 | IRF1 | JDP2 | ETV4 | |
